# Differential dominance of an allele of the Drosophila tßh gene challenges standard genetic techniques

**DOI:** 10.1101/504332

**Authors:** Christine Damrau, Julien Colomb, Björn Brembs

## Abstract

The biogenic amine octopamine (OA) and its precursor tyramine (TA) are involved in controlling a plethora of different physiological and behavioral processes. The *tyramine-ß-hydroxylase* (*tßh*) gene encodes the enzyme catalyzing the last synthesis step from TA to OA. Here, we report differential dominance (from recessive to overdominant) of the putative null *tßh^nM18^* allele in two behavioral measures in Buridan’s paradigm (walking speed and stripe deviation) and a proboscis extension assay in the fruit fly *Drosophila melanogaster*. The behavioral analysis of transgenic *tßh* expression experiments in mutant and wild type flies as well as of OA- and TA-receptor mutants revealed a complex interaction of both aminergic systems. Our analysis suggests that the different neuronal networks responsible for the three phenotypes show differential sensitivity to *tßh* gene expression levels. The evidence suggests that this sensitivity is brought about by a TA/OA opponent system modulating the involved neuronal circuits. This conclusion entails important implications for standard transgenic techniques, commonly used in functional genetics.

## Introduction

Pleiotropy is a central feature in genetics with pervasive implications for evolution [1–5]. Pleiotropic genes play an important evolutionary role not only because they create functional and developmental relationships among traits, but also because they can become relevant for the maintenance of genetic variability in a population [6,7]. While common, pleiotropy is not a universal property of all genes [8]. Pleiotropy is also a prerequisite for differential dominance. Differential dominance occurs when dominance patterns for a single locus vary among traits, e.g., the same allele may behave recessively in one trait and dominantly in another [4,9]. In wild populations, differential dominance is often accompanied by overdominance effects, thought to underlie the high level of heterozygosity found in these populations [4,6,7,9–11]. While heterozygosity tends to decrease in laboratory populations [12–18], the differential dominance effects may persist in pleiotropic genes, including overdominance.

Differential dominance, found to be ubiquitous in quantitative and population genetics studies [4,9,19], may potentially wreak havoc in functional genetics, where a common strategy is to introduce transgenic alleles into homozygously null mutant individuals [20–26]. For instance, if a mutation acts dominantly, rather than recessively, then the transgenic alleles will not rescue the phenotype even if the gene in question is responsible for it. In the case of overdominance, the outcome of such experiments may depend on the mechanism by which overdominance is achieved and could potentially range from no rescue to overdominant rescue, making these results difficult or impossible to interpret. Intermediate inheritance may make rescue experiments difficult to pin down statistically as successful or unsuccessful.

In the simplest case, the two alleles in question are a wildtype and a null mutant allele. In this arrangement, any differential dominance effects must be due to differential sensitivity of the phenotypes to gene dosage or gene expression levels or both. Therefore, such a situation is a good study case for investigating both the practical consequences for functional genetics studies and the mechanisms underlying the differential dominance phenomenon.

Because of the promiscuous role of biogenic amines in many different behavioral and physiological processes, the genes coding for their synthesis enzymes are prime candidates for pleiotropy and, hence differential dominance. The biogenic amine octopamine (OA) is structurally and functionally related to vertebrate noradrenaline [27–29]. OA is synthesized from another biogenic amine, tyramine (TA) by *tyramine-ß-hydroxylase* (*tßh*, [30]. OA plays an important role in the initiation and maintenance of motor programs in insects in general [31–35]. In skeletal muscles, OA concomitantly affects not only on the muscle tension [36] and relaxation rate [37], but also on its metabolism: As a neurohormone being released into the hemolymph, it mobilizes lipids and stimulates glycolysis [38,39]. OA appears to be involved in almost every behavioral and physiological process [40,41]. In Drosophila, the X-linked *tßh^nM18^* mutant has been an important tool to understand the role of TA and OA in many behaviors such as egg-laying [42–44], aggression [45,46], flight [47–50], and starvation resistance [51].

Loss-of-function *tßh^nM18^* male mutants, with a complete depletion of octopamine, display reduced aggression: their fight initiation latency is increased, while lunging and holding frequencies are decreased [46]. Furthermore, an acute silencing of octopaminergic neurons through the use of temperature-sensitive UAS-*Shi^ts^*, phenocopies the *tßh^nM18^* mutants, indicating the reduced aggression does not result from developmental defects in the mutants [45,52–54]. Interestingly, it was possible to rescue the aggression deficiency seen in *tßh^nM18^* mutant flies by expressing *tßh* in a small subset of octopaminergic neurons [45]. These results suggest that the standard genetic rescue approach can be successful at least in this phenotype, even with a pleiotropic gene. Some of us have shown previously that another phenotype, sugar sensitivity after starvation, can be analogously rescued [51].

In the present work we studied *tßh*-associated differential dominance and rescue experiments using behavioral phenotypes as disparate as sugar sensitivity [55,56] and walking behavior in Buridan’s paradigm [57,58]. In Buridan’s paradigm, we evaluated walking speed, a temporal parameter of movement control, as well as object fixation, a spatial measure of movement control. Fixation of visual cues is increased at higher contrast conditions [59,60]. Interestingly, the sensitivity of the motion-sensitive neurons in the fly optic lobes was shown to increase when the fly is walking [61–64] or flying [65]. The gain increase in flight was found to be OA dependent [66,67]. The three phenotypes we investigated (sugar responsiveness, walking speed and stripe fixation) exhibit differential dominance and we use various transgenic rescue techniques commonly used to elucidate gene function to probe the consequences of differential dominance on functional genetics studies as well as potential mechanisms mediating the differential dominance phenomenon. We complement these experiments with OA receptor manipulations in order to isolate OA-dependent from TA-dependent effects and to explore whether such gene-dosage-independent manipulations may be a superior functional genetic approach to mutant/rescue experiments.

## Materials and Methods

### Fly strains

*tßh^nM18^* (Monastirioti et al., 1996; FBal0061578), *oamb* ([68]; *OctαR*, *oamb286* FBti0038368, oamb584 FBti0038361), *honoka* (Kutsukake et al., 2000; Oct-TyrR, FBal0104701), *hsp*-*tßh* (Schwaerzel et al., 2003; FBal0152162), Octß2RΔ3.22 and Octß2RΔ4.3 (Damrau et al., 2014; CG6989, FBgn0038063), and w+;;UAS-*tßh* (Monastirioti, 2003; FBti0038601) were obtained from Henrike Scholz, Cologne; Hiromu Tanimoto, Martinsried; Andreas Thum, Konstanz; Martin Schwärzel, Berlin; and Amita Seghal, Chevy Chase. TyrR^f05682^ (CG7431f05682, FBal0184987), *TyrR^IIΔ29^* (CG16766, FBgn0038541) and *TyrR^II^-Tyr^RΔ124^* were kindly provided prior to publication by Edward Blumenthal, Milwaukee. Receptor mutants and their respective control lines were outcrossed for at least six generations into CS background.

**Table 1:**
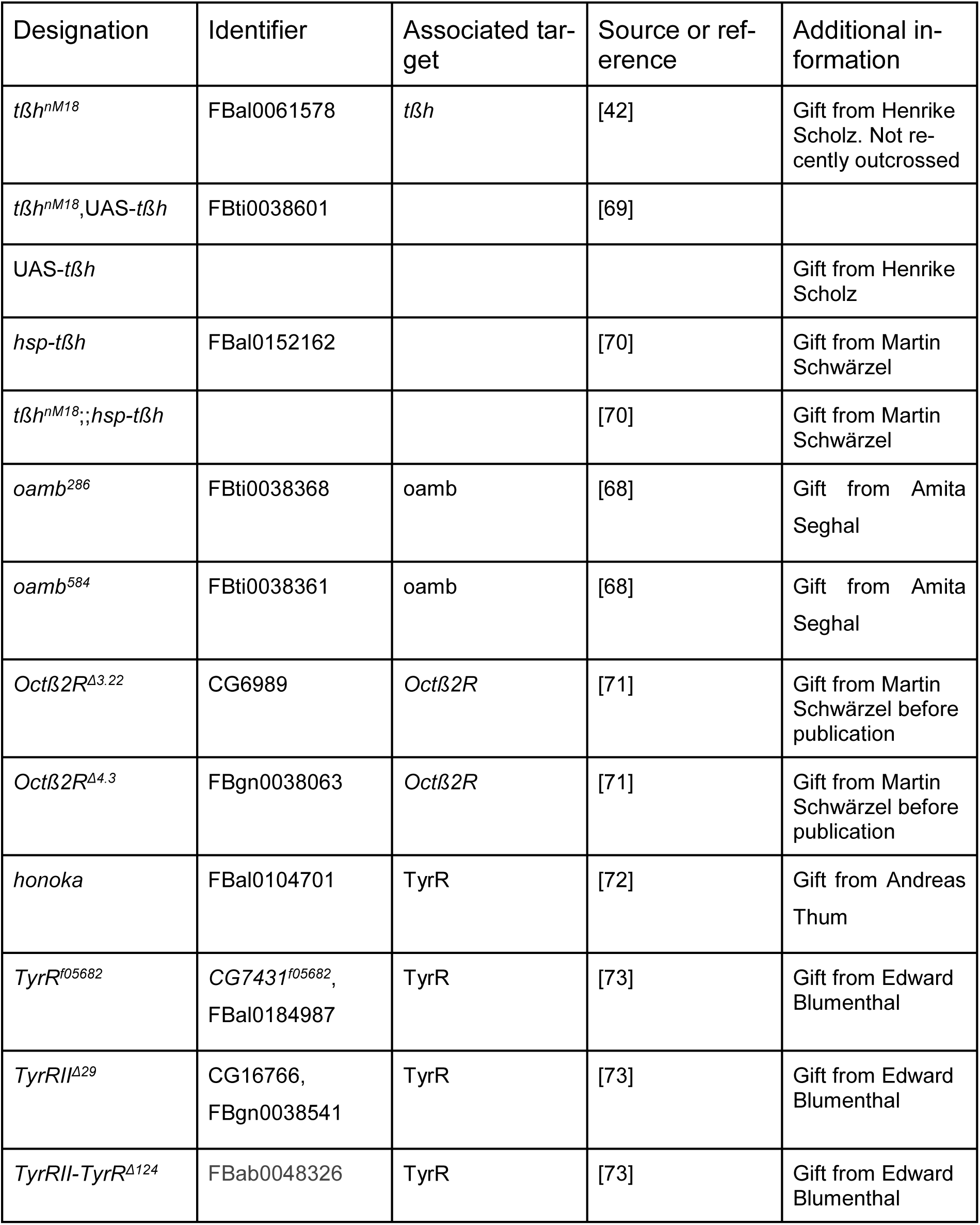
Fly strains used in this work.

The *tßh^nM18^* mutation is thought to be a null allele and abolishes OA synthesis. Consequently, the precursor of OA, TA, accumulates to approx. eight-fold over control levels [42]. Mutants for the Octß2R were created using recombination of FRT-containing P-elements (Parks et al., 2004), as described elsewhere [71]. In order to obtain hetero- and hemi-zygote mutants, we crossed *tßh^nM18^* mutant line with its original control line, which was also obtained from Henrike Scholz.

### Fly care

Flies were kept on standard cornmeal/molasses-food in a 12/12 h light/dark cycle at 60% relative humidity and 25°C except for hsp-*tßh* and elaV-GAL4;tub-GAL80 crosses which were kept at 18°C without humidity control.

After hatching, experimental flies were collected into new food vials for two days. The day before testing, flies were CO2-anesthetized, sorted by sex (females except for UAS-*tßh* experiments), and their wings were clipped at two thirds of their length. If not stated otherwise, animals recovered in the food vials overnight. Individuals were captured using a fly aspirator and transferred into the experimental setup on the following day.

### Heat-shock

hsp-*tßh* flies were heat shocked for 30-45 min at 37°C with 3-4 h recovery time at 25°C before testing. elaV-GAL4;tub-GAL80;UAS-Tdc2 flies were heated at 33°C overnight with 30 min recovery time at room temperature before testing.

### Buridan’s paradigm

We used the Buridan’s setup to test fly locomotion, details are described in [57](RRID:SCR_006331). Briefly, two black stripes (30 mm in width and 320 mm in height) were positioned opposite of each other 146.5 mm from the center of a platform (117 mm in diameter) surrounded by water and illuminated with bright white light from behind. The centroid position of the fly was recorded by custom tracking software (BuriTrack, http://buridan.sourceforge.net). If a fly jumped out of the platform, it was returned by a brush, and the tracker was restarted. All data are obtained from 5 min of uninterrupted walk or the first 5 min of a 15 min walk. See 10.17504/protocols.io.c7vzn5 for fly preparation.

Data were analyzed using CeTrAn v.4 (https://github.com/jcolomb/CeTrAn/releases/tag/v.4) as previously described in [57]. Briefly, walking speed was measured in traveled distance over time. A median was calculated for the progression of one experiment; the mean of all medians is reported in the graphs. Speeds exceeding 50 mm/s are considered to be jumps and are not included in the median speed calculation [57]. Stripe deviation acted as a metric for fixation behavior. It corresponds to the angle between the velocity vector and a vector pointing from the fly position towards the center of the frontal stripe (for details see [57]. Therefore, the larger the stripe deviation, the less accurate the fly fixated the stripe and vice versa. The platform inside the arena was cleaned with 70% ethanol after each experiment to minimize odor cues.

Buridan’s paradigm appears to be particularly sensitive to differences in genetic background [74]. Therefore, special emphasis was placed on always measuring all relevant genetic control lines simultaneously with the manipulated flies.

### Sugar sensitivity test

Sugar response was measured as described elsewhere [51]. Briefly, flies were starved for 20 h with Evian® water. Immobilized by cold-anesthesia using a cold station (Fryka-Kälteteschnik, Esslingen am Neckar, Germany), a triangle-shaped copper hook was glued to head and thorax. 3 h later, the hook was attached to a rack so that free movement of flies’ tarsi and proboscis was enabled. A filter paper soaked with sucrose solution was presented to all the tarsi. The proboscis extension response to a serial dilution of sucrose (0, 0.1%, 0.3%, 0.6%, 1%, 3% and 30%) was recorded. The total number of the fly’s responses to all sucrose stimulations of increasing concentration was calculated [75]. Finally, the proboscis was stimulated by 30% sucrose solution. Flies not responding to proboscis stimulation or responding to the first stimulation (water only) were discarded from the analysis.

### Statistics

The mean walking speed was calculated out of medians (see Buridan’s paradigm, above) and plotted with the standard error of the mean. Sucrose response and stripe deviation are shown as boxplots representing the median (bar), the 25%-75% quantiles, data within (whiskers) and outside (outliers as black dots) the 1.5 times interquartile range. Statistical analyses were performed in R (RRID:SCR_001905). Walking speed data followed normal distribution whereas stripe deviation did not (Shapiro-Wilk test of normality, p < 0.05) so that we used the parametric Two-way ANOVA followed by TukeyHSD post hoc test and Welch Two-Sample t-Test, respectively, or non-parametric paired Wilcoxon rank sum test with Bonferroni-correction and Wilcoxon rank sum test, respectively. The p-value was additionally corrected for two repeated measurements of data achieved from Buridan’s paradigm. The sample size of each group is indicated within the graphs. Default alpha value was set to 0.005 [76].

### Data availability

Raw data and evaluation code available at DOI: 10.5281/zenodo.4568550

## Results

### Differential dominance of *tßh* mutation for different behavioral parameters

We examined the effects of the *tßh^nM18^* mutation in two different experiments, assessing three different behavioral variables. We analyzed walking behavior in Buridan’s paradigm [57,58], reporting both the median speed (a temporal measure of behavior) and stripe deviation (as a spatial measure assessing object fixation). The second experiment quantified sugar responsiveness after 20 h of starvation using proboscis extensions [51].

Homozygous mutants behaved significantly differently from their genetic background-matched control flies in all three measures: homozygous female mutants showed reduced walking speed (Fig. 1A), fixated the stripes more closely (Fig. 1B), and were less likely to extend their proboscis to a sugar solution after starvation (Fig. 1C) compared to control flies with two intact *tßh* alleles. Flies heterozygous for the *tßh^nM18^* mutation did not behave similarly homogeneously across the three observed variables. In walking speed, heterozygous flies with only one intact *tßh* allele exceeded wildtype animals by about 20% (Fig. 1A), indicating overdominant inheritance. In stripe deviation, heterozygous flies behaved more similarly to the mutant flies than to the wild type control flies, indicating dominant inheritance (Fig. 1B). In the proboscis extension experiment, starved heterozygous mutant flies extended their proboscis two times (the median), compared to the one time for homozygous mutants and the three times for the wild type females (Fig. 1C). Despite being halfway between the two homozygous groups, indicating intermediate inheritance, the heterozygote data are not statistically different from the wild type controls, indicating a recessive phenotype.

**Fig. 1:**
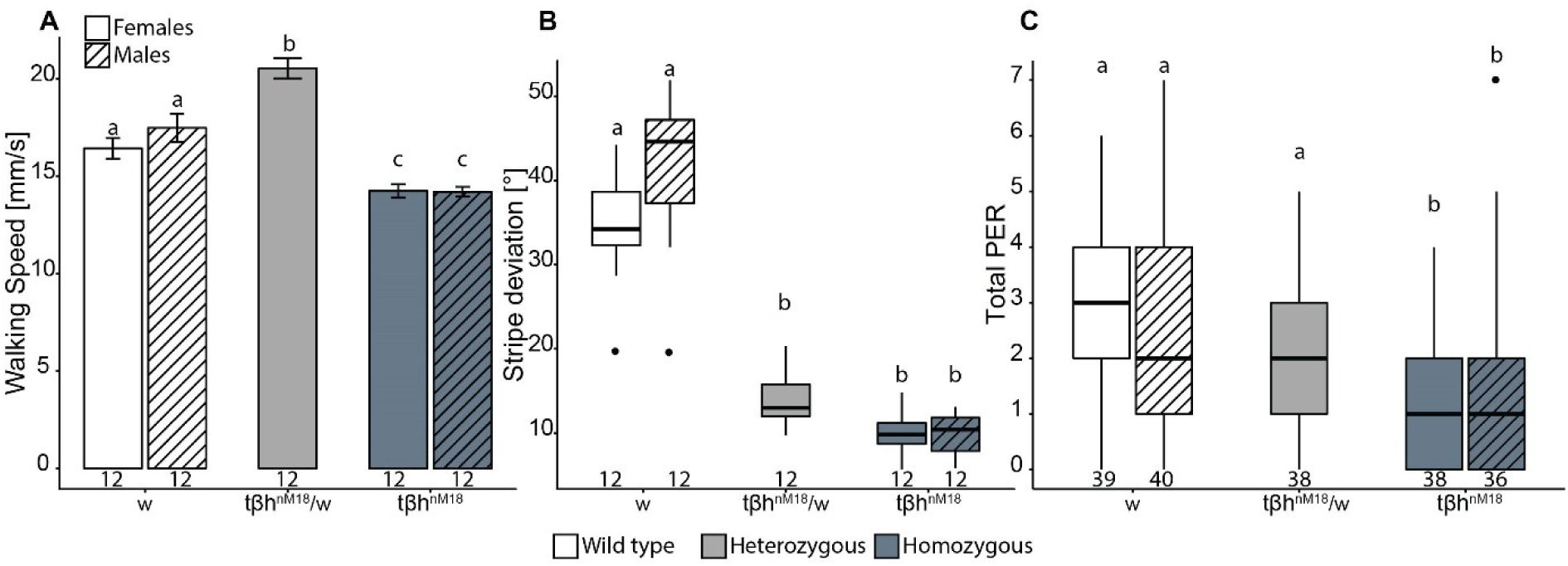
Differential dominance of *tßh* mutation for different phenotypes. Homo-, hetero- and hemizygous *tßh* mutants and their control were tested in Buridan’s paradigm and the sugar sensitivity test. **A.** Median walking speed in Buridan’s paradigm. Homozygous *tßh^nM18^* mutants walk slower than the wild type control, whereby heterozygous mutants walk faster than wild type (two-way ANOVA followed by TukeyHSD post hoc test, p < 0.005). Hemizygous mutant males walk slower than wild type males (Welch Two-Sample t-Test, p < 0.005). **B.** Stripe deviation, a measure of stripe fixation during walking, is reduced in homozygous *tßh^nM18^* mutants compared to heterozygous mutants and wild type (paired Wilcoxon rank sum test with bonferroni correction for repeated measurement, p < 0.005) as well as in hemizygous males compared to their wild type control (Wilcoxon rank sum test, p < 0.005). **C.** The total number of proboscis extension response to a serial dilution of sucrose after 20 h of starvation was calculated. Homozygous and hemizygous *tßh^nM18^* mutants respond less to sucrose compared to wild type controls; heterozygous mutants are not statistically different from wild type but different from homozygous mutants. Significant differences are tested by paired Wilcoxon rank sum test with Bonferroni correction (p < 0.005). In **A**, bars and error bars indicate mean and standard error of the mean. In **B** and **C**, the Tukey boxplots represent the median (bar), 25%-75% quartiles (boxes), and total data range (whiskers) excluding outliers outside of the 1.5x the interquartile range (points). Numbers below graphs indicate sample size. Bars and boxes labeled with different letters are statistically significantly different.

As some aspects of Buridan’s paradigm have been shown to be highly sensitive to genetic background [74] and the stripe deviation for the w+ control strain appeared unusually large, we repeated the locomotion experiment using flies with a different genetic background. We examined *tßh* hemizygous mutants and control males resulting from a cross to another wild type background. We found the same walking speed and stripe deviation phenotypes for *tßh* mutants in the W+/CantonS-background (p < 0.05, n = 34, Welch Two-Sample t-Test for speed, Wilcoxon rank sum test for stripe deviation; data at DOI: 10.5281/zenodo.2625643).

Taken together, we find that flies heterozygous for the *tßh^nM18^* mutation provide evidence for three different modes of inheritance, depending on the phenotype analyzed, conforming to the definition of differential dominance.

### Driving *tßh* expression via GAL4-UAS

What consequences can differential dominance have on standard genetic techniques, commonly leveraged to understand gene function? To tackle this question, we started with a tried-and-tested method of transgenically expressing a wild type version of the mutated gene in various tissues using the GAL4/UAS system [20–26,45]. As gene expression on the X-chromosome is doubled in hemizygous males, which behave indistinguishable from female flies (Fig. 1), the phenotypes of the heterozygous mutants (Fig. 1) may be due to the reduced expression of the single intact *tßh* gene. In other words, the failure of the heterozygous flies to behave like wild type flies may be due to the reduced gene dosage. To avoid such reduced gene expression in trans-heterozygous animals and to, instead mimic the hemizygous wild type males, we started our rescue experiments by driving an X-linked UAS-*tßh* transgene. We drove expression of the rescue construct in different tissues in *tßh* mutant males: in all cells (actin-GAL4), in all neurons (nSyb-GAL4), in tyraminergic non-neuronal cells (Tdc1-GAL4), in tyraminergic neurons (Tdc2-GAL4) and in octopaminergic neurons (NP7088-GAL4). All of those lines drive expression throughout development and in adulthood (Fig. 1). The X-linked transgene not only ensures doubled transcription from the single gene copy as in wild type males, it is also more practical as it is situated on the same chromosome as the mutation that is to be rescued. In fact, this chromosome was engineered precisely to make such rescue experiments more convenient than with the rescue transgene on an autosome, not unusual in functional genetics. Because we have already successfully used this technique on sugar responsiveness [51], we focus on the walking measures from now on. Neither the temporal nor the spatial walking measure showed any rescue for any of the targeted tissues (Fig. 2). In fact, for stripe deviation, some drivers yield even stronger stripe fixation than the mutant control strains (Fig. 2B). In both measures, some of the lines carrying the rescue construct alone already fail to show the mutant phenotype. Superimposed on the general trend of little effect in walking speed (Fig. 2A) and a reduction of stripe deviation (Fig. 2B), one can observe additional variability between the different groups. Presumably, this is due to the portions of differing genetic backgrounds the different GAL4 lines brought into the genotypes [74].

**Fig. 2:**
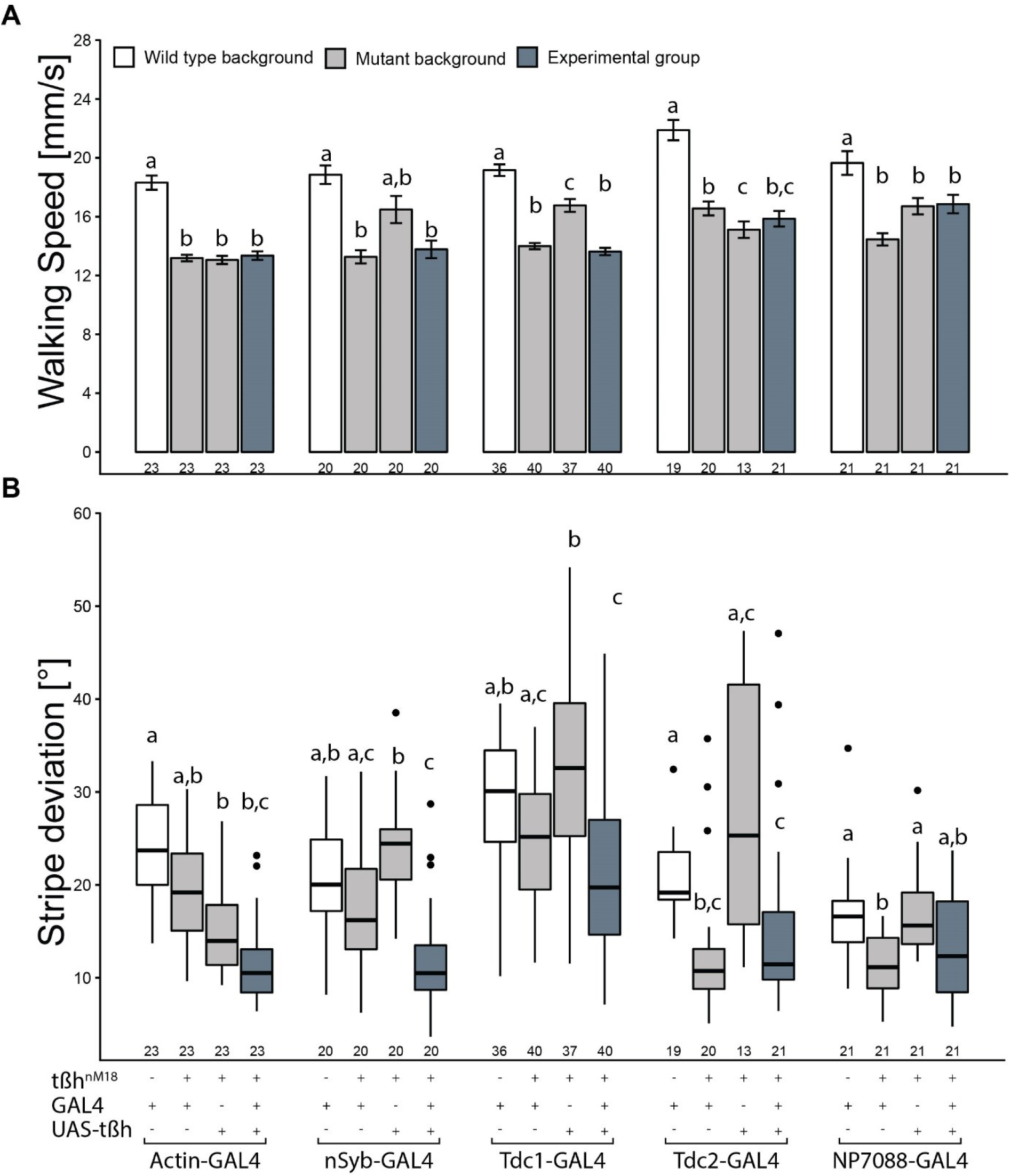
X-linked UAS-*tßh* expression cannot rescue the mutant Buridan phenotypes. **A**. Median walking speed cannot be rescued by heterozygous GAL4-UAS-dependent *tßh* expression in mutants. All groups are different from the wild type control, except for the UAS-*tßh* control in the nSyb experiment (two-way ANOVA with TukeyHSD post hoc test and correction for multiple measurements, p < 0.005). **B**. Stripe deviation performance is already increased by the presence of the GAL4- or the UAS-construct. Ubiquitous Actin-GAL4 or pan-neuronal nSyb-GAL4 expression worsens the phenotype compared to the control lines. Paired Wilcoxon rank sum test with Bonferroni correction, p < 0.005. Explanation for bars, Tukey box-plots, numbers and letters is identical to Fig. 1.

We had speculated that the heterozygous results (Fig. 1) may be due to low *tßh* transcription from the single gene dose. The rescue results (Fig. 2), on the other hand, may indicate that expression of too much *tßh* may also disrupt the walking behavior. To test this hypothesis, we performed the exact same experiments again, but this time with the UAS-*tßh* transgene on the third chromosome (Fig. 3), mimicking the situation in the heterozygous animals with halved gene expression, compared to the X-linked construct.

**Fig. 3:**
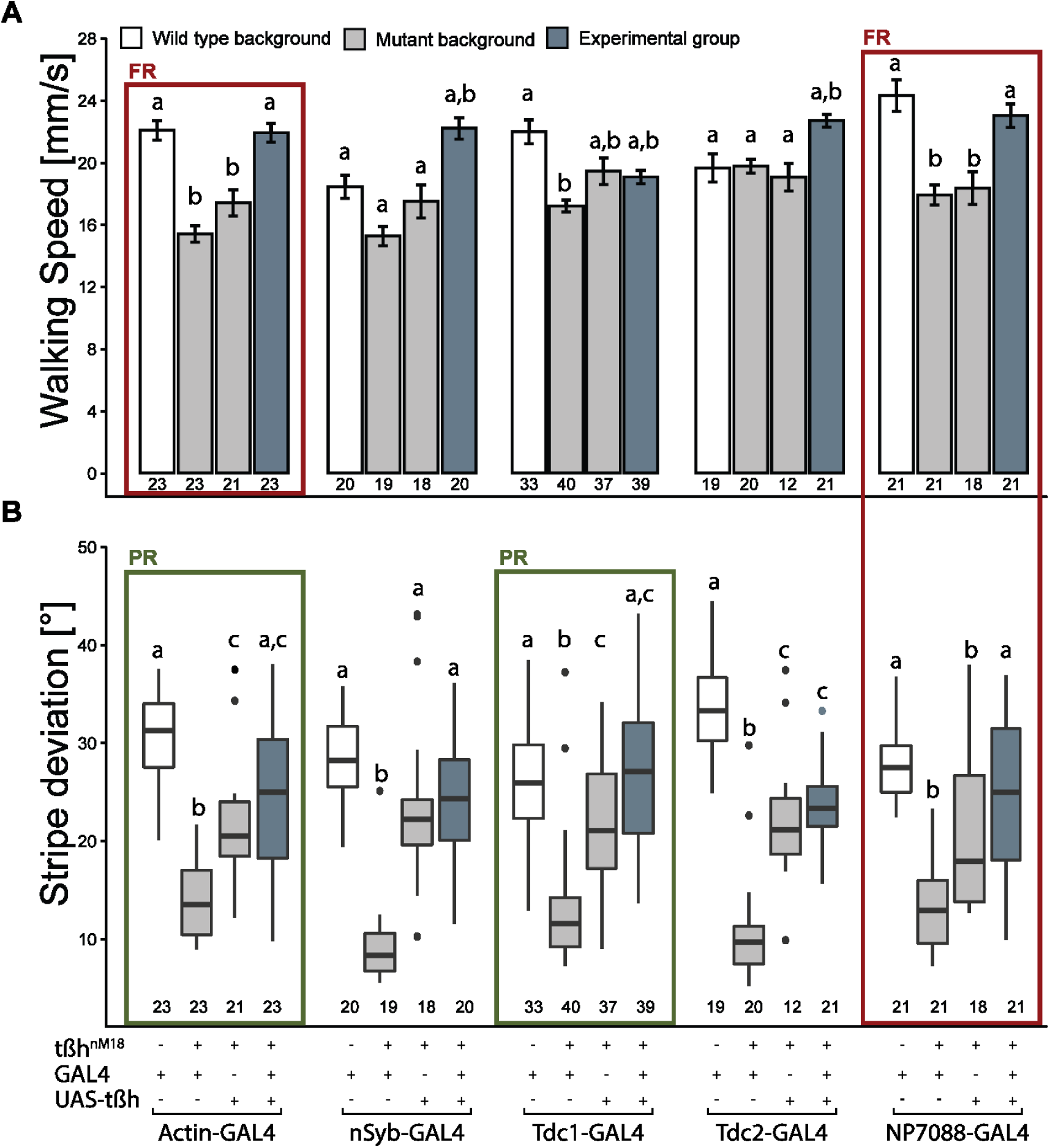
Autosomal UAS-*tßh* expression can rescue mutant Buridan phenotypes and phenocopy overdominance. **A**. Median walking speed can sometimes be rescued by heterozygous GAL4-UAS-dependent *tßh* expression in mutants. Only expression in all cells (actin-GAL4) and exclusively in octopaminergic cells via NP7088-GAL4 leads to a full rescue (FR, red) of the walking phenotype, characterized by significant differences of the experimental line (blue) from both mutant control groups (grey), but not from the wild type control (white). Two-way ANOVA with TukeyHSD post hoc test and correction for multiple measurements, p < 0.005. **B**. Stripe deviation performance is already increased by the presence of the UAS-construct. Only the octopaminergic NP7088-GAL4 rescues the stripe fixation phenotype. Expression of the transgene in all cells (Actin-GAL4) and in non-neuronal tyraminergic cells (TDC1-GAL4) leads to a partial rescue (PR, green), characterized by the experimental group failing to reach significant differences from either the wild type control or one of the mutant controls). Paired Wilcoxon rank sum test with Bonferroni correction, p < 0.005. Explanation for bars, Tukey boxplots, numbers and letters is identical to Fig. 1.

While again the mutant control strains with the rescue construct alone already showed some rescue effects, driving the rescue construct from the third chromosome yielded dramatically different results (Fig. 3) compared to the X-linked rescue attempts (Fig. 2). Octopaminergic (via NP7088-GAL4), but not tyraminergic (via Tdc-GAL4) expression led to a full rescue of both walking phenotypes, characterized by the rescue strain differing significantly from both mutant control strains but not from the wild type control. Expression in all cells (via Actin-GAL4) rescues walking speed completely, but the stripe fixation phenotype is only rescued partially. The overdominance in walking speed observed in heterozygous mutants was phenocopied in the pan-neuronal driver (nSyb-GAL4) as well as in the neuronal tyraminargic driver (Tdc2-GAL4). In both lines, the presence of the GAL4 constructs appears to already lower walking speed compared to the other wild type controls, making it indistinguishable from the mutant controls. Surprisingly, despite the expression pattern of Tdc2-GAL4 resembling that of NP7088-GAL4, there was no rescue of the stripe deviation phenotype, suggesting that stripe deviation is not influenced by different levels of tyramine in neurons. However, there was a partial rescue with the non-neuronal tyraminergic driver (Tdc1-GAL4), suggesting that non-neuronal tyraminergic cells (which do not express octopamine in wild type animals) influence stripe fixation.

These results suggest that the differential dominance of the *tßh* gene seems to be gene-dosage related. However, any deviation from wild type expression levels, both decreases and increases, can lead to significant differences to wild type behavior, rendering such standard experiments more of a lottery for pleiotropic genes that show differential dominance.

### Acute *tßh* expression differentially affects walking speed and stripe fixation

Another commonly used rescue technique is to ubiquitously express a wild type variant of the gene in the mutant background after development in the adult fly, i.e., right before the experiment. In our case, we expressed the *tßh* gene in homozygous *tßh^nM18^* mutant females under the control of the heat shock promoter hsp-*tßh*, situated on the third chromosome [70]. A heat shock was induced for 45 min at 37°C and flies were allowed to recover for 3 h. After this treatment, rescue flies walked faster than controls (Fig. 4A), phenocopying the overdominance results of the heterozygote flies (Fig. 1A). These results did not quite reach our stringent 0.005 alpha threshold, but passed the 0.05 threshold for suggestive effects. Given the behavior of the heterozygous flies, it is straightforward to assume an analogous over-dominance effect in this case. In contrast, expressing the *tßh* gene in this way left stripe fixation unaffected (Fig. 4B), similar to how heterozygous flies’ stripe deviation was indistinguishable that of homozygous mutant flies (Fig. 1B). As published previously [51], sugar response after heat shock rescue was significantly improved [Fig. 3C, from 51] without reaching wild type performance, similar to how heterozygous flies show an intermediate number of proboscis extensions when compared with the two homozygous groups (Fig. 1C).

**Fig. 4:**
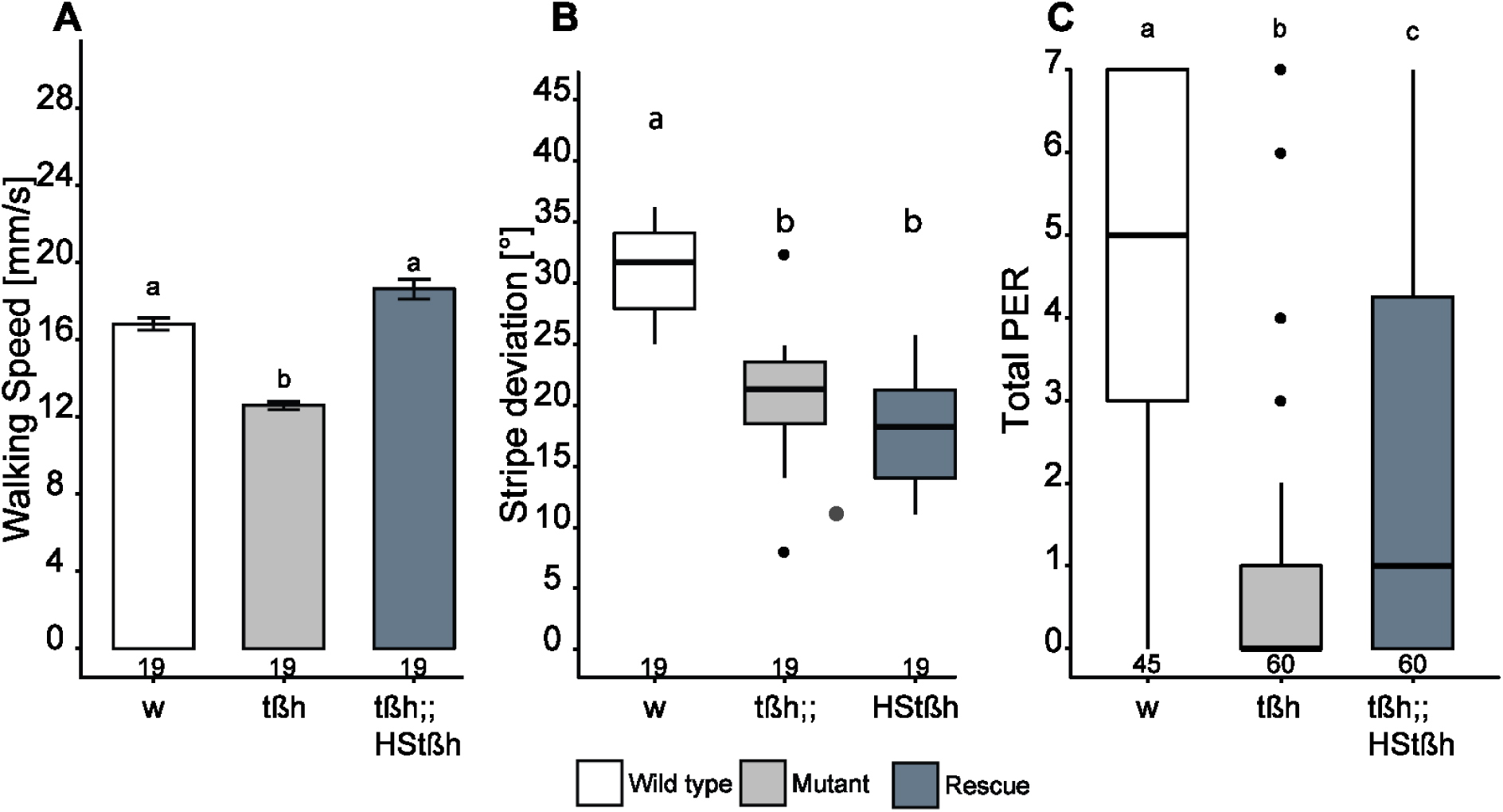
Acute and ubiquitous *tßh* expression rescues walking speed and sugar response but not stripe deviation. *tßh* expression was induced with a heat shock 3 h before the test. **A**. Median walking speed is increased beyond wild type levels after *tßh* induction (two-way ANOVA followed by TukeyHSD post hoc test, p < 0.005, F = 69.8). **B**. Stripe deviation is not affected by *tßh* induction. **C**. Sugar response is increased by *tßh* expression, but does not reach wild type levels (paired Wilcoxon rank sum test with Bonferroni correction, p < 0.005, data already published in Damrau et al. [51]). Explanation for bars, Tukey boxplots, numbers and letters is identical to Fig. 1.

Taken together, the results from heat shock-induced expression of *tßh* in the mutant background (Fig. 4) phenocopied those of the heterozygous flies (Fig. 1) throughout. Possibly, using a hsp-*tßh* construct on the X-chromosome may lead to a more successful rescue (i.e., opposite to the UAS rescue experiments). However, we are not aware of a *tßh^nM18^*,hsp-*tßh* X-chromosome.

### Overexpressing *Tdc2* and *tßh* differentially affects walking speed and stripe fixation

All experiments so far seem to suggest a very high sensitivity of the three chosen phenotypes to *tßh* gene dosage, where only a narrow range of gene expression supports wild type behavior. To test this hypothesis, we increased the acute expression of the *tßh* and Tdc (tyrosine decarboxylase; synthesizes TA from tyrosine) enzymes in wild type animals (Fig. 5). While *tßh* overexpression is assumed to lead to OA production from TA and hence a decrease in TA titers, the Tdc overexpression should lead to increased TA production and hence a subsequent increase in OA concentration as well.

**Fig. 5:**
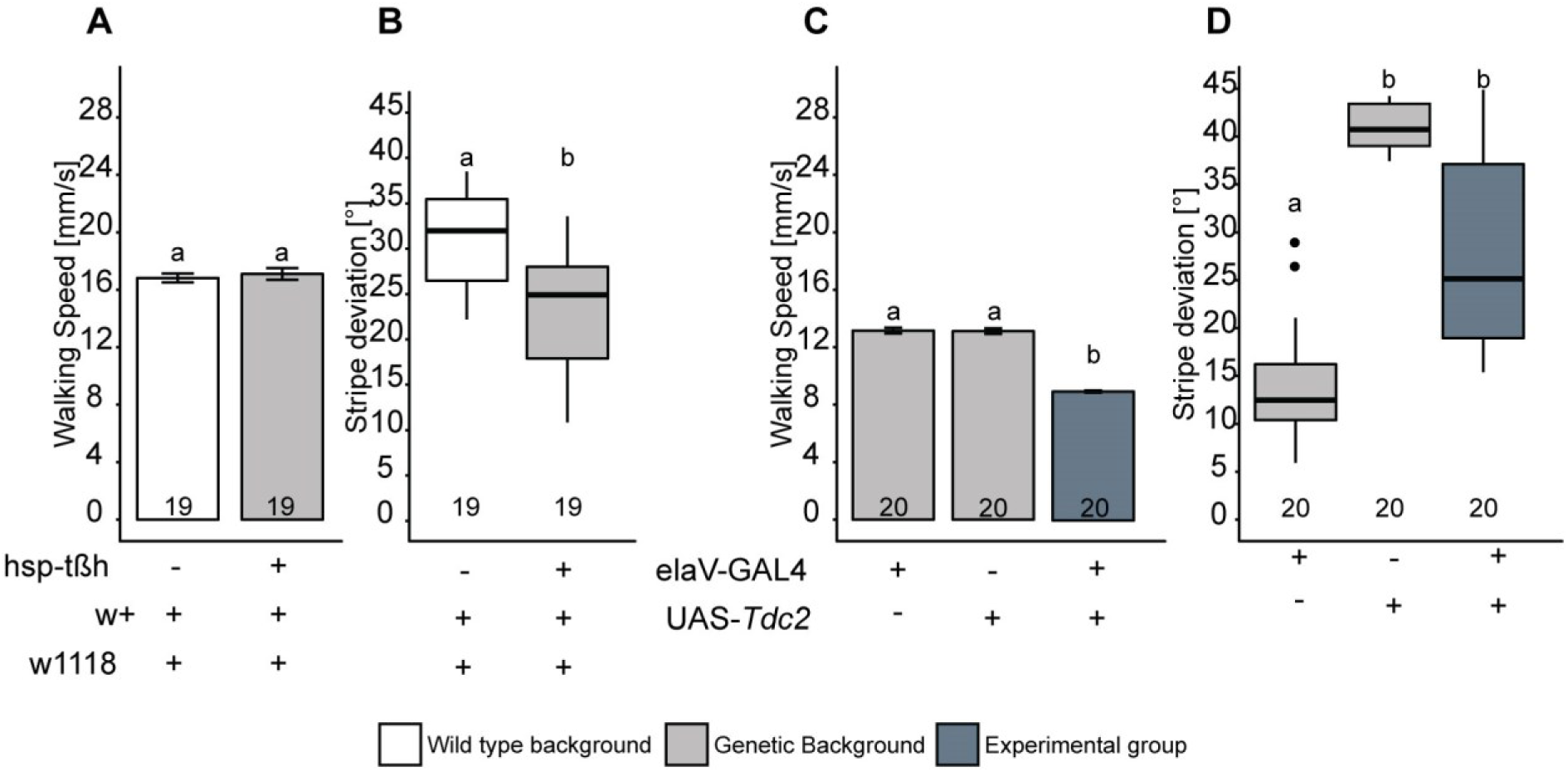
Acute overexpression of TA and OA synthesis enzymes in wild type background affects flies’ behavior in Buridan’s paradigm. **A**. Median walking speed is not affected when *tßh* is overexpressed in wild type 3 h before testing via *hsp-tßh* (Welch Two-Sample t-Test, p < 0.005). **B**. In contrast, stripe deviation is reduced when *tßh* is overexpressed (Wilcoxon rank sum test with correction for multiple measurements, p < 0.005). **C.** Median walking speed is reduced after overexpression of Tdc2 (two-way ANOVA followed by TukeyHSD post hoc test, p < 0.005). **D**. The two parental controls show very different stripe deviation behavior, which prevents the interpretation of the performance of the overexpression group (paired Wilcoxon rank sum test with Bonferroni correction, p < 0.005). Explanation for bars, Tukey boxplots, numbers and letters is identical to Fig. 1.

Overexpressing *tßh*, presumably decreasing TA levels and increasing OA levels, had no effect on walking speed (Fig. 5A), but decreased stripe deviation (Fig. 5B).Tdc2-overexpression, presumably elevating both TA and OA above wild type levels, reduced walking speed (Fig. 5B), but yielded a stripe deviation phenotype in the middle of the (large) range of variation found in driver and effector lines.

These results hence suggest that overexpressing *tßh* selectively affected the spatial measure stripe deviation, while overexpressing Tdc seemed to mainly affect the temporal measure walking speed. These results support the hypothesis that the Buridan phenotypes are exquisitely sensitive to *tßh* gene dosage. They also raise the possibility that the mechanism by which this sensitivity is achieved involves the relative levels of TA and OA, mediated by *tßh* expression. In order to investigate this possibility in a way that is both *tßh* gene dosage independent and can separately manipulate TA and OA signalling, respectively, we tested a number of TA and OA receptor mutants.

### Differential involvement of OA and TA receptors on walking speed and stripe fixation

To specifically affect the signaling of only one of the amines independently of the *tßh* locus, we manipulated the OA/TA system on the receptor level and examined several OA and TA receptor mutants, all outcrossed to the same genetic background (see methods).

We tested two alleles for each of two OA receptors *Oamb* and *Octß2R*, as well as one allele each for the three TA receptors *honoka*, *TyrR* and *TyrRII*, and a double receptor mutant for *TyrR* and *TyrRII*. While walking speed was affected in seven mutants (Fig. 6A, only TyrRIIΔ29 mutation had no effect), the stripe deviation was affected only in the two TA receptor mutants *TyrRf05682* and *honoka* in opposite directions (Fig. 6B). Interestingly, the double mutant *TyrRII*-*TyrRΔ124* showed no mutant phenotype in stripe fixation.

**Fig. 6:**
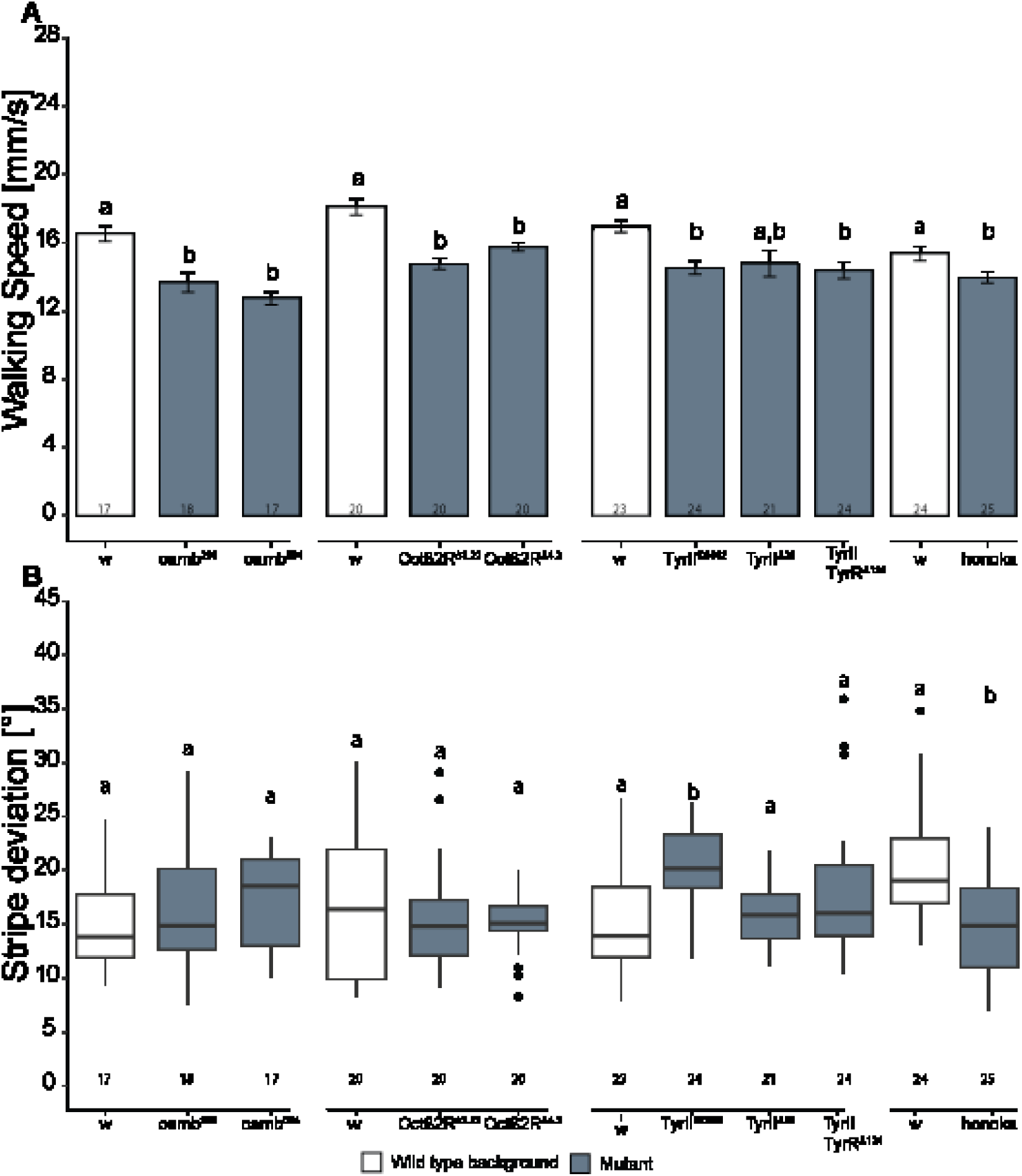
Performance of flies mutant for TA and OA receptors in the Buridan’s paradigm. **A**. *oamb^286^*, *oamb^584^*, *Octß2R^Δ3.22^*, *Octß2R^Δ4.3^*, *honoka*, *TyrR^f05682^*, and the double mutant *TyrRII-TyrR^Δ124^* walk slower than their respective controls, while *TyrRII^Δ29^* walking speed appeared at wild type level (Welch Two-Sample t-Test with correction for multiple measurements, p < 0.005). **B**. Stripe deviation is not affected in OA receptor mutants. In *TyrR^f05682^* and *honoka*, stripe deviation is significantly enhanced. Significant differences between control and the respective receptor mutant are calculated by Wilcoxon rank sum test with correction for multiple measurements (p < 0.005). Explanation for bars, Tukey boxplots, number and letters is identical to Fig. 1.

In principle, one would need to rescue each of the receptor mutants as well to exclude effects from genetic variations outside of the targeted locus. With regard to the results of the rescue experiments shown here, we have abstained from such experiments and rely, instead, on the extensive outcrossing of the lines to homogenize the genetic background between mutants and controls, to decrease the chances of off-target differences.

The results of the receptor experiments suggest OA being more specifically involved in walking speed, while TA signaling seems involved in both walking speed and stripe fixation. This latter hypothesis would be consistent with Tdc overexpression leading to reduced walking speed (Fig. 5B) and Tdc-1-GAL4 driven partial rescue of stripe fixation (in cells that do not express OA; Fig. 3B), but only if these phenotypes are also sensitive to deviation from wild type TA titers in both directions, increases and decreases.

## Discussion

### Differential dominance of the *tßh^nM18^* mutation

A likely null mutation [42] for the X-linked synthesis enzyme of the biogenic amine octopamine (OA), *tßh^nM18^*, showed differential dominance in three different behavioral traits (Fig.1). Pleiotropic alleles often show differential dominance [4,9] accompanied by overdominance in some of the traits [10,19]. However, it is not immediately obvious that the effects we observed are indeed attributable to differential dominance. Other phenomena that could lead to different outcomes for our different genotypes are dosage compensation in sex-linked genes (such as *tßh*), or other epigenetic effects, e.g., via differential maternal or paternal transfer of the gene in question (such as in our rescue experiments, Figs. 2, 3 and 4).

### Differential dominance is mediated by differential sensitivity to *tßh* gene expression levels

The *tßh* gene is located on the X-chromosome (X:7,995,697..8,027,394). The homozygous females and hemizygous males show identical phenotypes in all parameters tested, both in the mutant and the wildtype genotypes, respectively (Fig. 1). Thus, in these three cases, the presence of one or two X-Chromosomes appears to be irrelevant to the phenotype. Importantly, in males, the presence of one *tßh* allele is sufficient to provide the males with a wildtype phenotype for the parameters we studied. In the experiments where we tested heterozygous females (i.e., with only one intact allele of *tßh)* and find phenotypes indicating differential dominance, no dosage compensation takes place as this process occurs in males in *Drosophila* and not in females [77–82]. Presumably, the levels of *tßh* gene expression may be lower in heterozygous animals than in either wild type males or females. Importantly, this lower level of gene expression does not have the same effect in all phenotypes, leading to varying inheritance. Finally, the mutant allele in the heterozygous females always came from the mutant father, while a wild type mother (w+, with matched genetic background) provided the other X-Chromosome, such that any heterogeneity in the inheritance cannot come from heterogeneity in parent-of-origin imprinting effects, either. Thus, the only remaining explanation from the design of these experiments is that the pleiotropic *tßh* locus indeed confers differential dominance to the alleles we used here (Fig. 1), likely mediated by differential sensitivity of the three behaviors to *tßh* gene expression.

### Differential sensitivity to *tßh* gene expression levels is mediated by a TA/OA opponent system

Several studies have suggested that OA and TA may operate as an opponent system [47,83–86]. If this were the case, the raised levels of TA in the *tßh^nM18^* allele, may be partly responsible for some of the phenotypes observed, and not the lower OA levels. To our knowledge, it is still unknown if *tßh* is indeed the rate-limiting enzyme for OA synthesis or what effects manipulations of *tßh* expression levels may have on actual OA/TA titers. However, given that the OA precursor TA is also involved in locomotor control, it is straightforward to speculate that the acute sensitivity to *tßh* gene expression levels we have observed here may be reflecting a sensitivity to actual OA/TA titers in the neuronal networks involved. Thus, one potential mechanistic explanation for the differential dominance of the *tßh* locus is that the neuronal networks controlling the non-recessive behavioral parameters are modulated by a TA/OA opponent system which confers a high sensitivity to the relative amine titers (and hence gene expression levels) to network function.

Both the spatial rescue (Figs. 2, 3) and the overexpression results (Fig. 5) appear to support this hypothesis: we observed a partial rescue of stripe fixation in non-neuronal tyraminergic cells that do normally not release OA (Fig. 3B) and overexpressing the TA synthesizing enzyme *Tdc* affected walking speed.

To further explore this possibility, we manipulated TA and OA signaling individually, via OA receptor knock-out only in the non-recessive walking parameters (Fig. 6). In these experiments, stripe fixation measures a spatial property of walking behavior, as the flies need to orient themselves towards the stripes in space. Walking speed, in contrast, is taken as one of several measures of the temporal control of walking behavior in Buridan’s paradigm. These two parameters commonly separate not only in Principal Components Analyses, but also in biological manipulations.

#### Stripe fixation

All our X-linked GAL4/UAS manipulations of the *tßh* gene increased fixation behavior of the flies, even beyond wild type levels. Perhaps most strikingly, median stripe deviation was the lowest value for every single driver line we tested (Fig. 2B). This effect was also observed when driving gene expression with a heat-shock construct from the third chromosome (Fig. 4B). Only by presumably lowering the levels of gene expression by using a UAS construct on the third chromosome provided some successful rescue results. As part of such fixation behavior can be interpreted as an outcome of a fly’s light/dark preference [87], this dependence on gene expression levels may be understood by looking at photopreference results. Gorostiza et al. [87] discovered a correlation between dark-preference in a T-Maze and tighter fixation behavior in Buridan’s paradigm. With inhibited OA-ergic neurons, transgenic flies showed a lower dark-preference, while activated OA-ergic neurons increased dark preference. It is conceivable that both the doubled *tßh* gene expression from the dosage compensated X-chromosomes (Fig. 2B) and the hsp-driven rescue (Fig. 4B), increased the dark preference in these flies analogous to the activation of TA/OA-ergic neurons in [87].

Indeed, in our array of receptor mutants tested, OA receptor mutant flies do not fixate the stripes any different from control flies, while flies mutant for the TA receptors *honoka* and *TyrR^f05682^* fixate the stripes less strongly than wild type controls (Fig. 5B). In other words, decreased TA signaling can lead to decreased stripe fixation (Fig. 5B), while the increased TA levels in *tßh* mutants [42] can explain some of the increased fixation behavior in these flies (Fig. 1B). Although we have not tested all known OA receptors [88], considering the TA receptor and other data on stripe deviation, this suggests that TA may act independently of OA on this behavioral trait with an increased TA activity leading to stronger stripe fixation. We thus conclude that our manipulations of the OA synthesis enzyme *tßh* affected stripe fixation, at least in part, via an involvement of the OA precursor TA. This conclusion suggests that in *tßh^nM18^* mutants, the decreased stripe fixation may be due to the elevated levels of TA, while in our rescue and overexpression experiments, it may be due to high levels of OA, corroborating the hypothesis that TA/OA opponent organization may be the mechanism underlying the observed differential dominance effects.

#### Walking speed

The contrast to stripe fixation (a spatial measure of walking behavior) could not be starker in walking speed (a temporal measure of walking behavior). While it proved exceedingly difficult to decrease stripe fixation to control levels (observed in only one out of 13 manipulations), adding or removing *tßh* genes both increased and decreased walking speed. For instance, removing one copy (i.e., in the heterozygous state) increased walking speed, while removing both (homozygous mutants) decreased walking speed (Fig. 1A). Confirming the general observation that these two behavioral parameters are separable, also in these experiments walking speed and stripe fixation are decoupled.

Some lines driving transgenic expression of autosomal *tßh* rescue constructs (Fig. 2A), as well as acute *tßh* rescue before the experiment (Fig. 4A) yielded a phenocopy of the heterozygous flies: increased walking speed beyond wild type controls. At the same time, all X-linked rescue experiments failed to increase walking speed beyond mutant levels, suggesting that lower than normal *tßh* expression increases walking speed and higher than normal levels decrease it. This hypothesis is supported by our overexpression results: *Tdc2* over-expression throughout development reduces walking speed (Fig. 4A). These overexpression results in walking speed are mirror-symmetric compared to those in stripe fixation, supporting the potential opponent role OA/TA may be playing in both parameters and hence their role in establishing differential dominance in the *tßh* locus.

Due to these opposite results between spatial and temporal control of walking behavior and the highly varying nature of the walking speed results, one may speculate whether walking speed is either controlled by OA alone, or by OA in conjunction with TA. As we find that both OA and TA receptor mutants are affected in walking speed (Fig. 6A), we conclude that both OA and TA signaling are involved in the control of walking speed.

### Gene dosage in opponent systems

Manipulating *tßh* expression modifies the balance of OA and TA in opposite directions [42]. Therefore, the acute *tßh* gene expression dependence manifesting itself in differential dominance may be explained by the alteration of a fine balance between relative TA and OA concentrations. In *Drosophila* larvae, it was suggested that the relative increase in OA levels but not the absolute endogenous amount is important for regulation of starvation-induced locomotion [86]. However, the interaction between the two neuromodulators seems to be more complex than a simple balance [47,48,70,83–85,89–91]. We thus find that the data presented here are consistent with the hypothesis that one potential mechanism behind differential dominance in some traits is an opponent system of gene products which confers a high sensitivity to gene expression levels to these traits. These results support the hypothesis that the *tßh* gene exhibits type II pleiotropy [8].

### Differential dominance affects the outcomes of standard genetic techniques

As we have shown, this high sensitivity poses some formidable challenges for standard functional genetics techniques. A staple in the genetic toolbox are rescue experiments, which serve to establish the spatiotemporal expression requirements of the gene in question for the phenotypes under scrutiny [e.g., 23,24,45]. Such experiments are commonly carried out in order to arrive at necessity and sufficiency statements from which further mechanical understanding of gene function can follow [but see also 92,93]. However, the implicit and all too often untested assumptions for these experiments is that the (commonly) null mutations to be rescued follow recessive inheritance and that wild type level gene function can be restored with a single wild type allele. The GAL4/UAS system does not provide for sufficient control of gene expression levels to accommodate more unconventional modes of inheritance. In fact, in some cases, the basal promoter used in the creation of the GAL4 line may decide about the success or failure of an experiment [94].

In this work, we not only introduced the wild type allele of the *tßh* gene in its genomic locus in heterozygous animals (and hence with certainly wild type spatiotemporal expression levels; Fig. 1), but we also deployed commonly-used used spatial (Figs. 2, 3) and temporal (Fig. 4) transgenic rescue techniques, as well as transgenic overexpression in a wild type background (Fig. 5). While failed rescue experiments typically indicate that the mutated gene is not involved in the observed phenotype, the aggregate of all our experiments suggests that indeed the *tßh* gene is involved in all the phenotypes we studied, despite multiple failed rescue experiments in the walking phenotypes. Specifically, the autosomal or gonosomal location of the rescue construct affected rescue results via male dosage compensation (Figs. 2, 3), but the choice of technique driving the rescue construct, inasmuch as it affects expression levels, was also important irrespective of its autosomal location (Fig. 4). These data suggest that differential dominance can affect the outcome of some of these standard experiments to such an extent that nearly any arbitrary result may be obtained simply by the choice of rescue strategy - and the differential reporting of such results (file drawer effect) may distort the literature.

While pleiotropy was not found to be universal [8], it is not known how many genes in *Drosophila* are pleiotropic, nor how many of them display differential dominance. However, we have recently observed differential dominance in at least one more gene, the transcription factor *FoxP [94]*.

## Acknowledgements

We are grateful to the DFG for funding (BR 1892/9-1 in FOR1363), to Henrike Scholz, Hiromu Tanimoto, Andreas Thum and Amita Seghal for providing flies, to Edward Blumenthal for sharing data and flies prior to publication, to Martin Schwärzel and Sabrina Scholz-Kornehl for sharing their *Octß2R* mutants before publication, to Hildegard Hopp, Yasmine Graf, Julia Sigl, Lucie Dieterich, Luise Krüger and Sayani Banerjee for technical assistance, and to the workshop of Freie Universität Berlin for building the hardware.

## References

1. Fisher RA. The Genetical Theory of Natural Selection. Oxford University Press; 1930.

2. Orr HA. Adaptation and the cost of complexity. Evolution. 2000;54: 13–20.

3. Waxman D, Peck JR. Pleiotropy and the preservation of perfection. Science. 1998;279: 1210–1213.

4. Kenney-Hunt JP, Cheverud JM. Differential dominance of pleiotropic loci for mouse skeletal traits. Evolution. 2009;63: 1845–1851.

5. Otto SP. Two steps forward, one step back: the pleiotropic effects of favoured alleles. Proc Biol Sci. 2004;271: 705–714.

6. Turelli M, Barton NH. Polygenic variation maintained by balancing selection: pleiotropy, sex-dependent allelic effects and G x E interactions. Genetics. 2004;166: 1053–1079.

7. Lawson ND, Wolfe SA. Forward and reverse genetic approaches for the analysis of vertebrate development in the zebrafish. Dev Cell. 2011;21: 48–64.

8. Hill WG, Zhang X-S. On the pleiotropic structure of the genotype-phenotype map and the evolvability of complex organisms. Genetics. 2012;190: 1131–1137.

9. Ehrich TH, Vaughn TT, Koreishi SF, Linsey RB, Pletscher LS, Cheverud JM. Pleiotropic effects on mandibular morphology I. Developmental morphological integration and differential dominance. J Exp Zool B Mol Dev Evol. 2003;296: 58–79.

10. Klingenberg CP, Leamy LJ, Routman EJ, Cheverud JM. Genetic architecture of mandible shape in mice: effects of quantitative trait loci analyzed by geometric morphometrics. Genetics. 2001;157: 785–802.

11. Allison AC. Protection afforded by sickle-cell trait against subtertian malareal infection. Br Med J. 1954;1: 290–294.

12. Motro U, Thomson G. ON HETEROZYGOSITY AND THE EFFECTIVE SIZE OF POPULATIONS SUBJECT TO SIZE CHANGES. Evolution. 1982;36: 1059–1066.

13. Karlin S. Rates of Approach to Homozygosity for Finite Stochastic Models with Variable Population Size. Am Nat. 1968;102: 443–455.

14. Nowak C, Vogt C, Diogo JB, Schwenk K. Genetic impoverishment in laboratory cultures of the test organism Chironomus riparius. Environ Toxicol Chem. 2007;26: 1018–1022.

15. Saccheri I, Kuussaari M, Kankare M, Vikman P, Fortelius W, Hanski I. Inbreeding and extinction in a butterfly metapopulation. Nature. 1998;392: 491.

16. Madsen T, Shine R, Olsson M, Wittzell H. Restoration of an inbred adder population. Nature. 1999;402: 34.

17. Frankham R. Inbreeding and Extinction: A Threshold Effect. Conserv Biol. 1995;9: 792–799.

18. Dennis B. Allee effects in stochastic populations. Oikos. 2002;96: 389–401.

19. Cheverud JM, Ehrich TH, Vaughn TT, Koreishi SF, Linsey RB, Pletscher LS. Pleiotropic effects on mandibular morphology II: differential epistasis and genetic variation in morphological integration. J Exp Zool B Mol Dev Evol. 2004;302: 424–435.

20. Yokokura T, Dresnek D, Huseinovic N, Lisi S, Abdelwahid E, Bangs P, et al. Dissection of DIAP1 functional domains via a mutant replacement strategy. J Biol Chem. 2004;279: 52603–52612.

21. Chun Y-HP, Lu Y, Hu Y, Krebsbach PH, Yamada Y, Hu JC-C, et al. Transgenic rescue of enamel phenotype in Ambn null mice. J Dent Res. 2010;89: 1414–1420.

22. Kirshenbaum GS, Dachtler J, Roder JC, Clapcote SJ. Transgenic rescue of phenotypic deficits in a mouse model of alternating hemiplegia of childhood. Neurogenetics. 2016;17: 57–63.

23. Brand AH, Perrimon N. Targeted gene expression as a means of altering cell fates and generating dominant phenotypes. Development. 1993;118: 401–415.

24. Zars T, Fischer M, Schulz R, Heisenberg M. Localization of a short-term memory in Drosophila. Science. 2000;288: 672–675.

25. Saga Y. Genetic rescue of segmentation defect in MesP2-deficient mice by MesP1 gene replacement. Mech Dev. 1998;75: 53–66.

26. Park CJ, Zhao Z, Glidewell-Kenney C, Lazic M, Chambon P, Krust A, et al. Genetic rescue of nonclassical ERα signaling normalizes energy balance in obese Erα-null mutant mice. J Clin Invest. 2011;121: 604–612.

27. Evans PD. Biogenic Amines in the Insect Nervous System. In: Berridge MJ, Treherne JE, Wigglesworth VB, editors. Advances in Insect Physiology. Academic Press; 1980. pp. 317–473.

28. David J-C, Coulon J-F. Octopamine in invertebrates and vertebrates. A review. Prog Neurobiol. 1985;24: 141–185.

29. Roeder T. Octopamine in invertebrates. Prog Neurobiol. 1999;59: 533–561.

30. Wallace BG. The biosynthesis of octopamine-characterization of lobster tyramine β-hydroxylase. J Neurochem. 1976;26: 761–770.

31. Sombati S, Hoyle G. Central nervous sensitization and dishabituation of reflex action in an insect by the neuromodulator octopamine. J Neurobiol. 1984;15: 455–480.

32. Ridgel AL, Ritzmann RE. Insights into age-related locomotor declines from studies of insects. Ageing Res Rev. 2005;4: 23–39.

33. Baudoux S, Duch C, Morris OT. Coupling of efferent neuromodulatory neurons to rhythmical leg motor activity in the locust. J Neurophysiol. 1998;79: 361–370.

34. Duch C, Mentel T, Pflüger HJ. Distribution and activation of different types of octopaminergic DUM neurons in the locust. J Comp Neurol. 1999;403: 119–134.

35. Gal R, Libersat F. On predatory wasps and zombie cockroaches: Investigations of free will and spontaneous behavior in insects. Commun Integr Biol. 2010;3: 458–461.

36. Evans PD, O’Shea M. An octopaminergic neurone modulates neuromuscular transmission in the locust. Nature. 1977;270: 257–259.

37. Evans PD, Siegler MV. Octopamine mediated relaxation of maintained and catch tension in locust skeletal muscle. J Physiol. 1982;324: 93–112.

38. Candy DJ, Becker A, Wegener G. Coordination and Integration of Metabolism in Insect Flight*. Comp Biochem Physiol B Biochem Mol Biol. 1997;117: 497–512.

39. Blau C, Wegener G, Candy DJ. The effect of octopamine on the glycolytic activator fructose 2,6-bisphosphate in perfused locust flight muscle. Insect Biochem Mol Biol. 1994;24: 677–683.

40. Roeder T. Tyramine and octopamine: ruling behavior and metabolism. Annu Rev Entomol. 2005;50: 447–477.

41. Libersat F, Pflueger H-J. Monoamines and the Orchestration of Behavior. Bioscience. 2004;54: 17.

42. Monastirioti M, Linn CE Jr, White K. Characterization of Drosophila tyramine β-hydroxylase gene and isolation of mutant flies lacking octopamine. Journal of Neuroscience. 1996. Available: http://www.jneurosci.org/content/16/12/3900.short

43. Li Y, Fink C, El-Kholy S, Roeder T. The octopamine receptor octß2R is essential for ovulation and fertilization in the fruit fly Drosophila melanogaster. Arch Insect Biochem Physiol. 2015;88: 168–178.

44. Lim J, Sabandal PR, Fernandez A, Sabandal JM, Lee H-G, Evans P, et al. The octopamine receptor Octβ2R regulates ovulation in Drosophila melanogaster. PLoS One. 2014;9: e104441.

45. Zhou C, Rao Y. A subset of octopaminergic neurons are important for Drosophila aggression. Nat Neurosci. 2008;11: 1059–1067.

46. Baier A, Wittek B, Brembs B. Drosophila as a new model organism for the neurobiology of aggression? J Exp Biol. 2002;205: 1233–1240.

47. Brembs B, Christiansen F, Pflüger HJ, Duch C. Flight initiation and maintenance deficits in flies with genetically altered biogenic amine levels. J Neurosci. 2007;27: 11122–11131.

48. Pflüger H-J, Duch C. Dynamic neural control of insect muscle metabolism related to motor behavior. Physiology. 2011;26: 293–303.

49. Ryglewski S, Duch C, Altenhein B. Tyramine Actions on Drosophila Flight Behavior Are Affected by a Glial Dehydrogenase/Reductase. Front Syst Neurosci. 2017;11: 68.

50. O’Sullivan A, Lindsay T, Prudnikova A, Erdi B, Dickinson M, von Philipsborn AC. Multifunctional Wing Motor Control of Song and Flight. Curr Biol. 2018;28: 2705–2717.e4.

51. Damrau C, Toshima N, Tanimura T, Brembs B, Colomb J. Octopamine and Tyramine Contribute Separately to the Counter-Regulatory Response to Sugar Deficit in Drosophila. Front Syst Neurosci. 2018;11: 100.

52. Hoyer SC, Eckart A, Herrel A, Zars T, Fischer SA, Hardie SL, et al. Octopamine in male aggression of Drosophila. Curr Biol. 2008;18: 159–167.

53. Andrews JC, Fernández MP, Yu Q, Leary GP, Leung AKW, Kavanaugh MP, et al. Octopamine neuromodulation regulates Gr32a-linked aggression and courtship pathways in Drosophila males. PLoS Genet. 2014;10: e1004356.

54. Dierick HA. Fly fighting: octopamine modulates aggression. Current biology: CB. 2008. pp. R161–3.

55. Dethier VG. The hungry fly Harvard University Press. Cambridge, MA, USA. 1976.

56. Minnich DE. An experimental study of the tarsal chemoreceptors of two nymphalid butterflies. J Exp Zool. 1921;33: 172–203.

57. Colomb J, Reiter L, Blaszkiewicz J, Wessnitzer J, Brembs B. Open source tracking and analysis of adult Drosophila locomotion in Buridan’s paradigm with and without visual targets. PLoS One. 2012;7: e42247.

58. Bülthoff H, Götz KG, Herre M. Recurrent inversion of visual orientation in the walking fly,Drosophila melanogaster. J Comp Physiol. 1982;148: 471–481.

59. Osorio D, Srinivasan MV, Pinter RB. What causes edge fixation in walking flies? J Exp Biol. 1990;149: 281–292.

60. Riemensperger T, Isabel G, Coulom H, Neuser K, Seugnet L, Kume K, et al. Behavioral consequences of dopamine deficiency in the Drosophila central nervous system. Proc Natl Acad Sci U S A. 2011;108: 834–839.

61. Jung SN, Borst A, Haag J. Flight activity alters velocity tuning of fly motion-sensitive neurons. J Neurosci. 2011;31: 9231–9237.

62. Chiappe ME, Seelig JD, Reiser MB, Jayaraman V. Walking modulates speed sensitivity in Drosophila motion vision. Curr Biol. 2010;20: 1470–1475.

63. Rosner R, Egelhaaf M, Warzecha A-K. Behavioural state affects motion-sensitive neurones in the fly visual system. J Exp Biol. 2010;213: 331–338.

64. Maimon G, Straw AD, Dickinson MH. Active flight increases the gain of visual motion processing in Drosophila. Nat Neurosci. 2010;13: 393–399.

65. Li J, Lindemann JP, Egelhaaf M. Local motion adaptation enhances the representation of spatial structure at EMD arrays. PLoS Comput Biol. 2017;13: e1005919.

66. Suver MP, Mamiya A, Dickinson MH. Octopamine neurons mediate flight-induced modulation of visual processing in Drosophila. Curr Biol. 2012;22: 2294–2302.

67. Strother JA, Wu S-T, Rogers EM, Eliason JLM, Wong AM, Nern A, et al. Behavioral state modulates the ON visual motion pathway of Drosophila. Proc Natl Acad Sci U S A. 2018;115: E102–E111.

68. Han K-A, Millar NS, Davis RL. A Novel Octopamine Receptor with Preferential Expression inDrosophila Mushroom Bodies. J Neurosci. 1998;18: 3650–3658.

69. Monastirioti M. Distinct octopamine cell population residing in the CNS abdominal ganglion controls ovulation in Drosophila melanogaster. Dev Biol. 2003;264: 38–49.

70. Schwaerzel M, Monastirioti M, Scholz H, Friggi-Grelin F, Birman S, Heisenberg M. Dopamine and octopamine differentiate between aversive and appetitive olfactory memories in Drosophila. J Neurosci. 2003;23: 10495–10502.

71. Scholz-Kornehl S. Generierung und Charakterisierung der Deletionsmutanten des G-Protein gekoppelten Rezeptors D2R von Drosophila melanogaster. 2015 [cited 21 Dec 2018]. Available: https://refubium.fu-berlin.de/handle/fub188/5978

72. Kutsukake M, Komatsu A, Yamamoto D, Ishiwa-Chigusa S. A tyramine receptor gene mutation causes a defective olfactory behavior in Drosophila melanogaster. Gene. 2000;245: 31–42.

73. Zhang H, Blumenthal EM. Identification of multiple functional receptors for tyramine on an insect secretory epithelium. Sci Rep. 2017;7: 168.

74. Colomb J, Brembs B. Sub-strains of Drosophila Canton-S differ markedly in their locomotor behavior. F1000Res. 2014;3: 176.

75. Scheiner R, Page RE, Erber J. Sucrose responsiveness and behavioral plasticity in honey bees (Apis mellifera). Apidologie. 2004;35: 133–142.

76. Benjamin DJ, Berger JO, Johannesson M, Nosek BA, Wagenmakers E-J, Berk R, et al. Redefine statistical significance. Nat Hum Behav. 2018;2: 6–10.

77. Lucchesi JC, Kuroda MI. Dosage compensation in Drosophila. Cold Spring Harb Perspect Biol. 2015;7. doi:10.1101/cshperspect.a019398

78. Lucchesi JC. Transcriptional modulation of entire chromosomes: dosage compensation. J Genet. 2018;97: 357–364.

79. Moschall R, Gaik M, Medenbach J. Promiscuity in post-transcriptional control of gene expression: Drosophila sex-lethal and its regulatory partnerships. FEBS Lett. 2017;591: 1471–1488.

80. Birchler JA. Parallel Universes for Models of X Chromosome Dosage Compensation in Drosophila: A Review. Cytogenet Genome Res. 2016;148: 52–67.

81. Baker BS, Gorman M, Marín I. Dosage compensation in Drosophila. Annu Rev Genet. 1994;28: 491–521.

82. Conrad T, Akhtar A. Dosage compensation in Drosophila melanogaster: epigenetic fine-tuning of chromosome-wide transcription. Nat Rev Genet. 2012;13: 123–134.

83. Saraswati S, Fox LE, Soll DR, Wu C-F. Tyramine and octopamine have opposite effects on the locomotion of Drosophila larvae. J Neurobiol. 2004;58: 425–441.

84. Fox LE, Soll DR, Wu CF. Coordination and modulation of locomotion pattern generators in Drosophila larvae: effects of altered biogenic amine levels by the tyramine β hydroxlyase mutation. Journal of Neuroscience. 2006. Available: http://www.jneurosci.org/content/26/5/1486.short

85. Alkema MJ, Hunter-Ensor M, Ringstad N, Horvitz HR. Tyramine Functions independently of octopamine in the Caenorhabditis elegans nervous system. Neuron. 2005;46: 247–260.

86. Koon AC, Budnik V. Inhibitory control of synaptic and behavioral plasticity by octopaminergic signaling. J Neurosci. 2012;32: 6312–6322.

87. Gorostiza EA, Colomb J, Brembs B. A decision underlies phototaxis in an insect. Open Biol. 2016;6. doi:10.1098/rsob.160229

88. Farooqui T. Review of octopamine in insect nervous systems. Open access insect physiol. 2012. Available: https://www.researchgate.net/profile/Tahira_Farooqui/publication/269598262_Review_of_octopamine_in_insect_nervous_system/links/549819280cf2c5a7e342960c.pdf

89. Scheiner R, Plückhahn S, Oney B, Blenau W, Erber J. Behavioural pharmacology of octopamine, tyramine and dopamine in honey bees. Behav Brain Res. 2002;136: 545–553.

90. Fussnecker BL, Smith BH, Mustard JA. Octopamine and tyramine influence the behavioral profile of locomotor activity in the honey bee (Apis mellifera). J Insect Physiol. 2006;52: 1083–1092.

91. Hoyle G. Generation of Behaviour: the Orchestration Hypothesis. Feedback and Motor Control in Invertebrates and Vertebrates. 1985. pp. 57–75.

92. Yoshihara M, Yoshihara M. “Necessary and sufficient” in biology is not necessarily necessary - confusions and erroneous conclusions resulting from misapplied logic in the field of biology, especially neuroscience. J Neurogenet. 2018;32: 53–64.

93. Gomez-Marin A. Causal Circuit Explanations of Behavior: Are Necessity and Sufficiency Necessary and Sufficient? In: Çelik A, Wernet MF, editors. Decoding Neural Circuit Structure and Function. Cham: Springer International Publishing; 2017. pp. 283–306.

94. Palazzo O, Rass M, Brembs B. Identification of circuits involved in locomotion and object fixation in. Open Biol. 2020;10: 200295.

